# Are populations of economically important bonefish and queen conch ‘open’ or ‘closed’ in the northern Caribbean Basin?

**DOI:** 10.1101/2020.07.08.194415

**Authors:** Marlis R. Douglas, Tyler K. Chafin, Julie E. Claussen, David P. Philipp, Michael E. Douglas

## Abstract

Demographics of co-occurring species can often be diagnosed through population genomic analyses of single nucleotide polymorphisms (SNPs). These data can define population structure, gene flow, and candidate regions in the genome that potentially reflect local adaptations. They can also gauge if populations are demographically ‘open’ or ‘closed’ (i.e., with global or local recruitment). We derived SNPs from double-digest restriction-site associated DNA (ddRAD) to test the demographics of commercially important bonefish *Albula vulpes* (N=117) and queen conch *Lobatus gigas* (N=60) from two northeast Caribbean Basin islands (Grand Bahama to the north and Eleuthera to the south). Specifically, we tested the hypothesis that the strong west-to-east current in the Great Bahama Canyon is a vicariant barrier separating the two islands. We conducted Bayesian assignment tests on putatively neutral (*A. vulpes=*36,206 SNPs; *L. gigas*=64,863) and highly-differentiated outlier datasets (*A. vulpes=*123 and 79 SNPs; *L. gigas*=88 and 51, respectively). For bonefish, results diagnosed asymmetrical gene flow and north-south differentiation, as potentially driven by adult mobility and easterly currents. However, both analyses indicated genetic structure in conch, substantiating the vicariant hypothesis. These results provide templates for future research endeavors with these impacted species. Outlier loci, for example, can potentially place populations of each within a demographic continuum, rather than within a dichotomous ‘open/closed’ framework, as well as diagnose ‘source’ and ‘sink’ populations, as herein. These methodologies can then be applied to co-distributed species with similar but less well understood ecologies so as to evaluate basin-wide trends in connectivity and local adaptation.

## 1 INTRODUCTION

Connectivity among populations is a fundamental consideration for both marine conservation (Bell et al. 2014) and management (Miller et al. 2014), with aspects that resonate both theoretically (Kool et al. 2010, Ellingson & Krug 2015) and functionally (Hameed et al. 2016, Krueck et al. 2017). Benthic marine invertebrates and many sedentary fishes generally disperse as a component of the plankton so as to reach areas beneficial for growth and development. This often extends across temporally variable periods with the end result being populations diagnosed as demographically ‘open’ (Domingues et al. 2010). Many marine fishes follow a similar life-history strategy (Williamson et al. 2016), but with an additional and innate capacity to ‘self-recruit’ back into their natal populations (Christie et al. 2010), a situation that confounds the traditional ‘open’ versus ‘closed’ dynamic. In a similar vein, larger-bodied marine vertebrates often exhibit philopatry, defined as the return of reproducing individuals to the locality from which they were spawned (Feldheim et al. 2014, Bonanomi et al. 2016, Salles et al. 2016). This, in turn, necessitates a capacity to home with great precision, in that natal areas were previously experienced only during the egg or larval stage. When this behavior is manifested in both sexes, a ‘closed’ population can result such that dynamics are driven not by long-distance immigration but rather by local reproduction and recruitment (Bentzen & Bradbury 2016).

Numerous mechanisms have been employed to evaluate dispersal and connectivity in marine species, and consequently, to diagnose their open or closed status (Cowen et al. 2000, Cowen & Sponaugle 2009). In this regard, much contemporary evidence stems from indirect methods, with particular emphasis on genetic and genomic techniques that gauge ‘effective dispersal’ (i.e., that due to survival and subsequent reproduction), rather than direct observation of individual movements among populations. The former provide considerably greater resolution, and often establish larval relatedness (Ottmann et al. 2016), self-recruitment (Teske et al. 2016), panmixia (Jasonowicz et al. 2017, Kensington et al. 2017, Tay et al. 2017), and adult dispersal (Veríssimo et al. 2017). Genomic assays have also been extended as well to estimate dispersal and connectivity among deep-sea biodiversity, where studies are largely dominated by more conservative methods (Baco et al. 2016).

While methods of dispersal in marine biodiversity are many and varied, their variance can often be a source of ambiguity, particularly when statistical tests of revelant hypotheses are conducted (Weersing & Toonen 2009). For example, several studies have displayed varying results when population connectivity was tested against the temporal duration of the planktonic larval stage. Some recognized a positive correlation between connectivity and distance (Reisser et al. 2014, Ellingson & Krug 2015, Herrera et al. 2016, Morvezen et al. 2016, Tay et al. 2016, Van Wyngaarden et al. 2016, Jasonowicz et al. 2017, Kensington et al. 2017, Martinez et al. 2017). Yet, others have demonstrated an inverse relationship (Kool et al. 2010, Jackson et al. 2014, Jorde et al. 2015, Cumming et al. 2016, Gomes et al. 2016, Hameed et al. 2016, Teske et al. 2016). Clearly, dispersal can be constrained or accelerated by many factors, with the promotion or depression of connectivity being an end result.

In a similar vein, other studies have established global interconnectedness but, contrastingly, as a complex matrix of locally-adapted populations, with larval transport varying both spatially and temporally (Christie et al. 2010, Miller et al. 2014, Gould et al. 2017, McKeon et al. 2017, Segovia et al. 2017). This, in turn, suggests a gradient in population structure, rather than a binary condition where available choices are either ‘open’ *versus* ‘closed.’ Here, we investigated the existence of binary spatial population structure as potential endpoints of a continuum, and did so by examining populations of economically important marine biodiversity in the northeastern Caribbean Basin. Our focus on a distant corner of the Caribbean ecosystem was based on the presence of an extensive and deep marine trench (the Great Bahama Canyon) that may serve as a potential west-to-east vicariant barrier that prevents larval gene flow between populations of bonefish *Albula vulpes* and queen conch *Lobatus gigas* located at islands to the north and south, respectively.

### 1.1 Bonefish

A complex of four bonefish species occur within the Atlantic Ocean/ Caribbean Basin: Bonefish (*Albula vulpes*), Threadfin bonefish (*A. nemoptera*), Channel bonefish (*A. goreensis*), and an as yet undescribed form *A. sp. cf. vulpes*. The three nominate species are partitioned non-geographically in the Basin (Wallace & Tringali 2016). We restricted our evaluation to *A. vulpes*, a species with an elevated home range fidelity within the shallow flats proximate to Bahamian islands (Murchie et al. 2013). However, deeper channels separaing shallow flats can serve as conduits for travel or as refugia from rapid changes in water temperature (Humston et al. 2005).

Inshore/nearshore habitats are somewhat distant from home ranges, yet allow pre-spawning bonefish to congregate during October-May, such as on the south side of Grand Bahama island (Figure 1) where deeper spawning waters are available (Murchie et al. 2015). Spawning movements into offshore waters then occur on the full moon (Danylchuk et al. 2011), with adults gathering at depth (>130m) prior to a quick ascent (~67m in <2min) to spawn within a distinct oceanographic layer of low turbulence and flow (i.e., optimal pycnocline/ thermocline) that subsequently retains the planktonic leptocephali larvae (Lombardo et al. 2020). The latter subsequently drift for ~41–71 days before eventually recruiting to unspecified inshore nurseries (Adams & Cook 2015).

**FIGURE 1.**
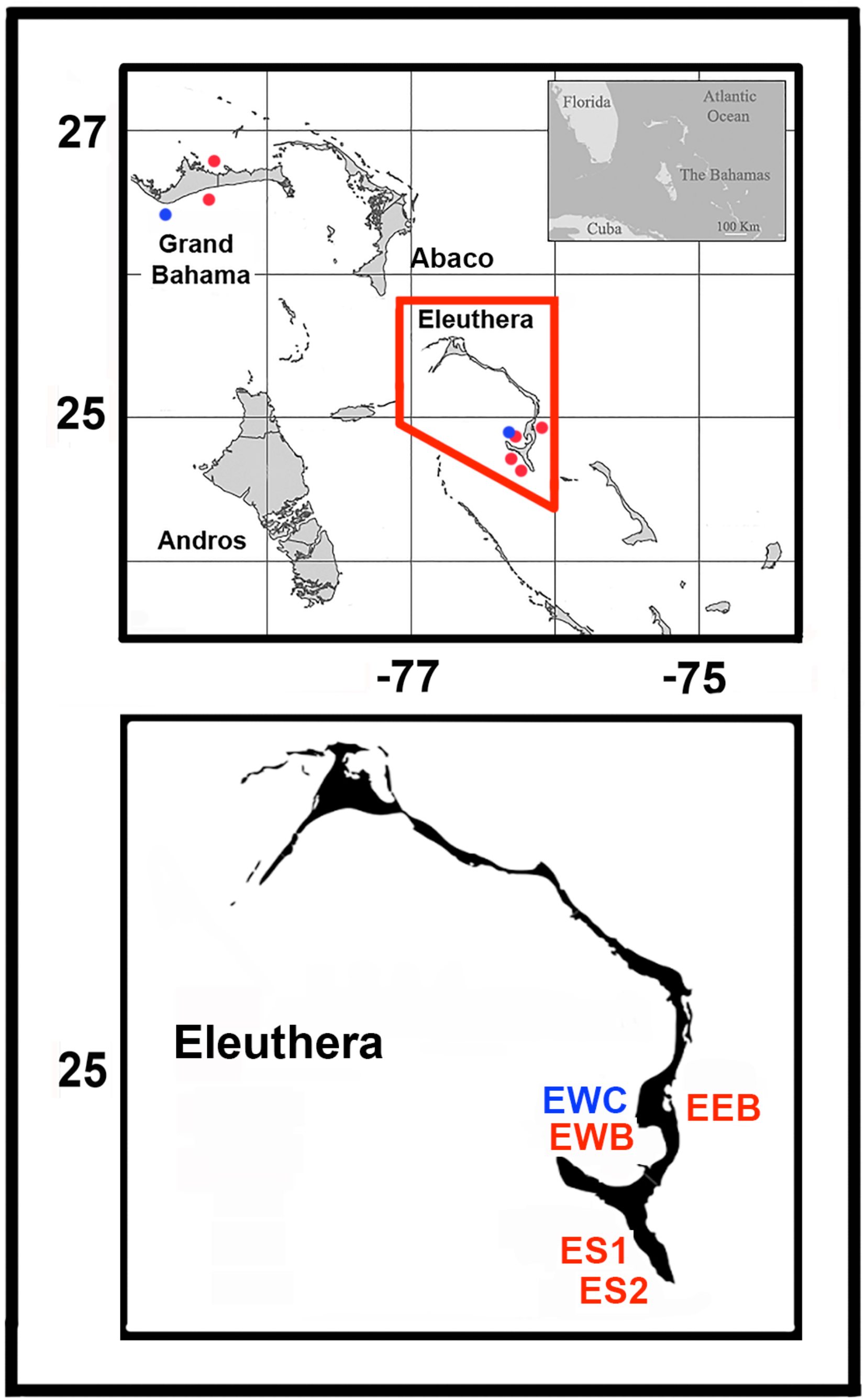
(Top) Map depicting The Commonwealth of Bahama (insert top right). The islands of Grand Bahama, Abaco, Eleuthera and Andros (northeastern Basin) are depicted. Blue dots represent sample sites for queen conch, whereas those for bonefish are designated with red dots. (Bottom) Sample sites at Eleuthera Island for queen conch and bonefish. Colors as above. Labels as in Table 1.

A basin-wide mtDNA analysis of *A. vulpes* (Wallace & Tringali 2016) revealed two somewhat intermixed genetic populations. However, the extent of their larval drift from local or distant sources is unknown. Populations of *A. vulpes* may reflect a situation found in other high gene flow species, such as the cardinalfish *Siphamia tubifer*, where genetic homogeneity effectively conceals a high population turnover and a variable accumulation of larvae from divergent sources (Gould et al. 2017). The brown crab, *Cancer pagurus,* displays a similar condition in the English Channel and North Sea, where high levels of dispersal and gene flow effectively mask a genetic patchiness that results from a recruitment that is spatially and temporally variable (McKeon et al. 2017).

**TABLE 1.** Collection data for bonefish (*Albula vulpes*) and queen conch (*Lobatus gigas*) at Grand Bahama and Eleuthera islands, northern Caribbean Basin. Listed are: Species, Island, Side (of island with compass direction in parentheses), Locality, Date, N (=sample size), and Label (=sample abbreviation).

| Species | Island | Side | Locality | Date | N | Label |
| --- | --- | --- | --- | --- | --- | --- |
| Bonefish | Grand Bahama | Flats (N) | North | 26-Jan-14 | 30 | GNB |
| Bonefish | Grand Bahama | Ocean (S) | South | 25-Jan-14 | 30 | GSB |
| Bonefish | Eleuthera | Flats (W) | Broad Creek | 2-Mar-14 | 30 | EWB |
| Bonefish | Eleuthera | Ocean (E) | Half Sound | 8-Mar-14 | 10 | EEB |
| Bonefish | Eleuthera | Flats (SW) | Plum Creek | 8-Mar-14 | 10 | ES1 |
| Bonefish | Eleuthera | Flats (SW) | Sound Side | 10-Mar-14 | 7 | ES2 |
| Conch | Grand Bahama | South | Freeport | 1-Jan-14 | 30 | GSC |
| Conch | Eleuthera | Flats (W) | Schooner Cay 2 | 1-Mar-14 | 30 | EWC |

### 1.2 Queen conch

Queen conch (*Lobatus gigas*) is not only an icon of considerable economic and cultural importance (Kough et al. 2017), but also represents the largest of six species in the Caribbean Basin. It feeds on micro-and macro-algae, but with habitat shifts that occur ontogenetically from shallower seagrass beds into deeper-water macroalgal plains (~40m in depth) (Doerr & Hill 2013). Harvest (and over-harvest) are promoted by a density-dependent reproductive strategy in which both sexes not only reduce movements but also aggregate along open sandy bottoms for broadcast spawning (Stoner 1997).

Spawning extends from April through October, with peak larval abundance in late June-August when temperatures in the northeastern Caribbean Basin exceed 28°C (Stoner & Davis 1997a). Planktonic larvae drift for ~60–75 days (Ballantine & Appeldoorn 1983), but with high natural mortality and with distances controlled by oceanic conditions. For example, the surface environment dictates the vertical distribution of larvae in the water column, with >80% of larvae occurring at 0.5-5.0m depth (Stoner & Davis 1997b).

In 1992, *L. gigas* was recognized by the Convention on International Trade of Endangered Species (CITES: https://cites.org/eng/prog/queen_conch/introduction) and was subsequently added in 1994 to the Red List maintained by the International Union for the Conservation of Nature (IUCN; http://www.fao.org/docrep/006/Y5261E/y5261e07.htm). A review of the *L. gigas* fishery is provided by Brownell & Stevely (1981).

A broad-ranging spatial genetic study of *L. gigas* across the greater Caribbean using nine microsatellite DNA markers (Truelove et al. 2017a,b) found that basin-wide gene flow was constrained by oceanic distance that served to isolate local populations, but surprisingly, not within Bahamian populations where *F*_ST_ values dis not differ significantly. Similarly, a more regional study in the Mesoamerican Caribbean using five inter-microsatellite DNA (ISSR) primers again rejected panmixia (Machkour-M’Rabet et al. 2017), whereas an earlier protein electrophoretic study in the Bahamas indicated gene flow was more homogeneous (Mitton et al. 1983). Also of note is the fact that overall genetic homogeneity among populations (per the Bahamas) can be masked by the annual variation in numbers of larvae produced by over-fished source populations.

### 1.3 Research objectives

In this study, we employed a double-digest restriction-site associated DNA (ddRAD) methodology to identify single nucleotide polymorphisms (SNPs) as diagnostic markers in bonefish and conch. Our study is the first to employ this techniques with these economically important study species, and we hypothesized that our derivation of neutral loci would reflect the homogenizing effects of gene flow in each, thus emphasizing their placement within the ‘open’ end of the demographic spectrum. We then sought to contrast our initial results employing neutral loci by deriving a more reduced and fine-grained series of outlier SNPs within each species that may potentially demonstrate local adaptation. By doing so, we tested the hypothesis that sharp environmental gradients within a vicariant marine topography can counterbalance gene flow and promote instead adaptive divergence.

Finally, we recognize that our study species comprise exploited stocks, each of which transects numerous national borders across the Caribbean Basin. As such, they experience varying levels of temporal and spatial exploitation. Thus, a more nuanced understanding of coarse- and fine-grained dispersal in these species, and indeed if they display connectivity, provides a necessary baseline from which to derive an ecologically meaningful framework for sustainable conservation. In this sense, the potential for local adaptation and source-sink dynamics between islands could provide a rationale from which to seriously recalibrate conch and bonefish management in the Caribbean.

## 2 MATERIALS AND METHODS

### 2.1 Study site

The Caribbean Basin extends westward from Florida along the Gulf Coast of North America, then south along the coast of México and Central America, continuing eastward across the northern coast of South America, with end-points at the Greater and Lesser Antilles. Within this matrix, the Commonwealth of the Bahamas (Figure 1 top) represents a shallow series of carbonate banks (~104,000km^2^) situated on a submerged continental border east of Florida (Andrews et al. 1970). Our Bahamian study sites are located along the coasts of two islands: Grand Bahama (153km west-east; 24km north-south), and Eleuthera (~1.6km west-east, 180km north-south; Figure 1 top). Grand Bahama is located 86km east of Palm Beach (FL) and represents the fourth largest island in the Bahamian chain, whereas Eleuthera is some 370km distant, with the Atlantic Ocean on its east side and the shallow (<3m depth) Great Bahama Bank immediately west (Figure 2 top). Approximately 1–3km southwest lie the abyssal waters of Exuma Sound (~1500m depth; “3” of Figure 2 top).

**FIGURE 2.**
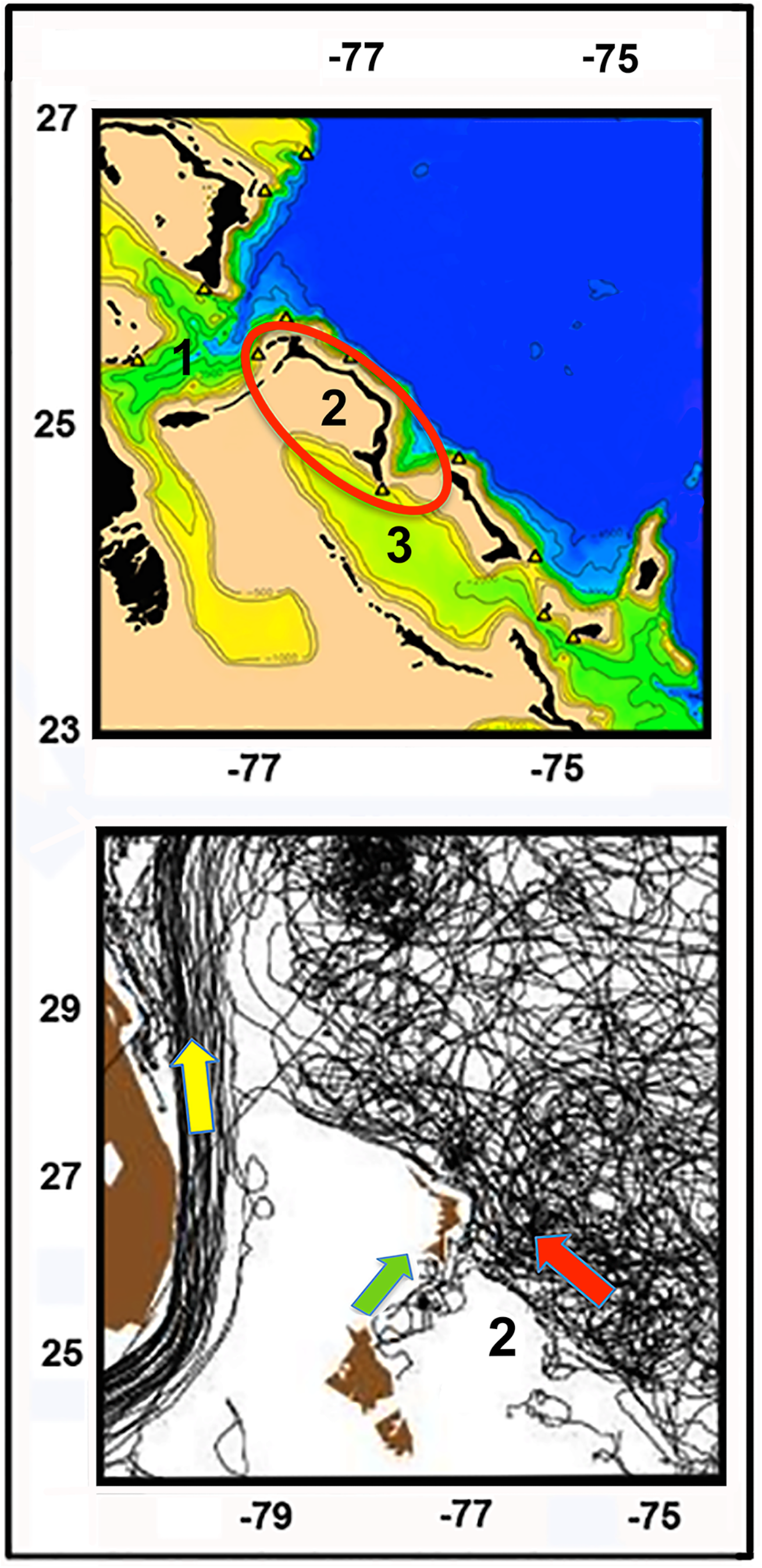
(Top) Bathymetry map of the northwest Carribean Basin (modified from http://oceanexplorer.noaa.gov/explorations/09deepseacorals/logs/summary/media/bahamas_region_600.html). Colors indicate depth: Blues = deepest (>3,000m); yellow-to-green = intermediate (500-2500m); beige = shallowest (<500m); Islands depicted in black. Red ellipse = Eleuthera Island; Great Bahama Canyon = 1; Great Bahama Flat = 2; Exuma Sound = 3. (Bottom) Oceanic currents in the northern Caribbean Basin (modified from: University of Miami Rosenthiel School of Marine and Atmospheric Science, http://oceancurrents.rsmas.miami.edu/caribbean/caribbean-maps.html). Black lines = currents; Yellow arrow = direction of Gulf Stream (east coast of Florida); Green arrow = current direction in Grand Bahama Canyon; Eleuthera Island = 2; Red Arrow = direction of Antillean Current (Atlantic Ocean).

The Great Bahama Canyon (length=225km, max width=37km, max depth=6.9km;“1” of Figure 2 top) is located northeast of the Great Bahama Bank, and effectively separates Grand Bahama and Abaco islands (to the north) from Eleuthera (to the south; Figure 1). It is divided into two main branches: ‘Northwest Providence Channel’ extending north and west towards Grand Bahama Island, with ‘Tongue of the Ocean’ extending south along the east coast of Andros Island, facing Exuma Sound. The Northwest Providence Channel also absorbs a small but significant easterly contribution from the Gulf Stream (yellow arrow, Figure 2 bottom; Richardson & Finlen 1967), with currents extending west-to-east in the Canyon (green arrow, Figure 2 bottom). The occasionally turbid currents within the Great Bahama Canyon deposit a large sediment fan as they merge with the Antilles Current of the Atlantic Ocean ~65km to the north-northeast (red arrow, Figure 2 bottom) (Andrews 1970). Although the seasonal variability of these currents has not been established, their overall direction and extent offer the potential for a vicariant separation between Grand Bahamas and Eleuthera.

### 2.4 Collection, extraction, and amplification of samples

During 2014, we sampled fin clips from 57 adult bonefish collected by rod and reel from three locations on the west and south sides of Eleuthera Island (EWB, ES1, ES2; Table 1) and one from the Atlantic Ocean (east) side (EEB; Table 1). Muscle samples were also gathered from 60 adults at Grand Bahama Island, with 30 taken from the south (Ocean) side (GSB) and 30 from the north (Flats) side (GNB) (total N=117). Sampling was in compliance with the U.S. National Research Council Guide for the Care and Use of Laboratory Animals (https://grants.nih.gov/grants/olaw/Guide-for-the-Care-and-use-of-laboratory-animals.pdf).

We also collected 30 individuals of queen conch from the south side of Grand Bahama Island (GSC), and 30 from the Flats (or west) side of Eleuthera Island (EWC) (Table 1, Figure 1 bottom). Fin clips and muscle plugs were preserved on Whatman^®^ FTA^®^ Classic DNA cards, with genomic DNA subsequently extracted using a Gentra^®^ Puregene DNA Purification Kit and stored in hydrating solution (per manufacturer instructions). The quality of DNA was visualized on 2% agarose gels, then quantified using a Qubit fluorometer (Thermo-Fisher Scientific, Inc.). Subsequent library preparation followed previously published protocols (Peterson et al. 2012, as modifed in Bangs et al. 2018).

Restriction digests were performed using 1μg of genomic DNA with *Pst*I (5’-CTGCAG-3’) and *Msp*I (5’-CCGG-3’) in CutSmart buffer (New England Biosciences) for 20 hours at 37°C, after first optimizing restriction enzyme and size selection choices using an *in silico* approach (FRAGMATIC; Chafin et al. 2018). Digests were visualized on 2% agarose gels, purified using AMPure XP beads, and quantified with a Qubit fluorometer. Digested samples were normalized to 0.1μg of DNA and subsequently ligated with barcoded Illumina adaptors, using custom oligos (Peterson et al. 2012). Inline barcodes were designed to differ by at least 2 bases so as to minimize mis-assignments. Pooled ligations were size-selected at 350-400bps using the Pippin Prep automated size fractionator (Sage Sciences).

Adaptors were extended by PCR (10 cycles) with Phusion high-fideltiy DNA polymerase (New England Biosciences). Size distribution and successful amplification of purified libraries was then confirmed on the Agilent 2200 TapeStation. Indexed libraries were multiplexed with 100-bp single-end sequencing on the Illumina HiSeq 2500 lane (Genomics and Cell Characterization Core Facility, University of Oregon/Eugene).

### 2.5 Processing and assembly of ddRAD data

Raw ~100bp reads were demultiplexed and processed in the PYRAD pipeline (Eaton 2014) using a 90% identity threshold for clustering homologous reads. Read clusters representing putative loci within individuals were excluded when sequencing depth was <10. Following assembly, additional loci were discarded if they exhibited: 1) <50% presence across individuals; or 2) >10 consensus N bases. We additionally removed potential paralogs or over-merged loci by excluding those having: 1) >2 alleles in any individual; 2) >60% heterozygosity per site among individuals; 3) >10 heterozygous sites per consensus sequence. Further data processing was performed using custom scripts (available at: https://github.com/tkchafin/scripts).

### 2.6 Detection and spatial structure of local selection

We first sub-sampled all SNPs to a maximum of one per locus. We then utilized a Bayesian method (BAYESCAN v.2.1, Foll & Gaggiotti 2008) with a burn-in period of 50,000 followed by 100,000 iterations to generate a candidate list of potential loci under selection. We then used *F*_ST_-etimates to derive three additional datasets per taxon: (1) All unlinked SNPs (=ALL); (2) SNPs >2 standard deviations (sd) above mean *F*_ST_ (=SD2); and (3) SNPs >3 sd above the mean (=SD3).

We employed multiple methods to examine population structure for both species across all three datasets. Hierarchical iterative clustering analysis (Structure v.2.2.4; Pritchard et al. 2000) was run in parallel (Besnier & Glover 2013) on the Jetstream cloud computing platform (Hancock et al. 2019) using the GNU Parallel shell tool (Tange 2011). Structure was run for 250,000 repetitions with a burn-in period of 100,000, and with potential clusters for K-values ranging from 1-20. Optimal K for each dataset was ascertained using the delta-K method (Evanno et al. 2005), as implemented and visualized in the CLUMPAK pipeline (Kopelman et al. 2015). Differentiation among sampling localities was also examined with with a cross-validation approach (Discriminant Analysis of Principal Components, DAPC) in the R package adegenet (Jombart 2008, Jombart et al. 2008). We defined the number of retained principal component axes as that which minimized the root-mean-square error (RMSE) of assignment for 10% of randomly selected individuals (with the remaining 90% serving as a training set), and N=30 replicates per level of PC retention. We specified the number of DF axes retained as equal to the number of populations minus one.

### 2.7 Asymmetric migration among populations

Physical processes such as water current and tidal fluxes often create patterns of asymmetric gene flow among populations, and consequently impact patterns of genetic diversity. Given our initial hypothesis, we were interested in gauging the amount and direction of gene flow among the six study populations of bonefish, and two of queen conch. In contrast to Structure, which employs a Bayesian probabilistic model to assign individuals to ancestry clusters, we instead estimated the posterior probability of individual migratory histories, thereby allowing the direction and magnitude of dispersal among populations to be dissected.

To do so, we employed a revised software optimized for large SNP datasets (i.e., BA3-SNPS, Mussmann et al. 2019). We first tuned MCMC acceptance parameters using an automated iterative procedure, then generated posterior estimates of migration rates using 250,000 iterations with a burn-in period of 50,000. We further tested the hypothesis of source-sink dynamics by computing net emigration rates as a sum of immigration rates subtracted from summed emigration rates (Andreasen et al 2012), with positive numbers indicating a genetic source population whereas negative values indicated a sink population.

## 3 RESULTS

After filtering, we obtained 56,046 loci for bonefish *Albula vulpes*, of which 68.7% (N=36,206) were parsimoniously informative (Table 2). Average number of loci per individual was 35,443, at a mean coverage depth of 77x. Subsampling only unlinked SNPs produced a matrix of 25,205 SNPs (=ALL), from which datasets SD2 (123 SNPs) and SD3 (79 SNPs) were derived, respectively.

For queen conch *Lobatus gigas*, 30,697 loci were available post-filtering, and represented 64,864 SNPs, of which 68.4% (N=44,334) were parsimony informative. Individuals were sequenced at a coverage depth of 58.6x, averaging 33,409 loci per individual (Table 2). Subsampled unlinked SNPs totaled 19,716 (=ALL), from which datasets SD2 (88 SNPs) and SD3 (51 SNPs) were obtained.

**TABLE 2.** Study species (= bonefish *Albules vulpes*; queen conch *Lobatus gigas*) evaluated in this study. Also provided are number of genetic loci greater than 50% retention (= Loci > 50%); Average number of loci per individual (= Av loci/ind); Average depth of coverage (= Av coverage); Number of parsimonious-informative SNPs (= PI SNPs); Number of unlinked SNPs (=ALL); and number of SNPs sampled at 2 (=SD2) and 3 (=SD3) standard deviations from the mean *F*_ST_.

| <u>Species</u> | <u>Loci > 50%</u> | <u>Av loci/ind</u> | <u>Av coverage</u> | <u>PI SNPs</u> | <u>ALL</u> | <u>SD2</u> | <u>SD3</u> |
| --- | --- | --- | --- | --- | --- | --- | --- |
| <u>Bonefish</u> | <u>56,046</u> | <u>35,443</u> | <u>77X</u> | <u>36,206</u> | <u>25,205</u> | <u>123</u> | <u>79</u> |
| <u>Conch</u> | <u>30,697</u> | <u>33,409</u> | <u>59X</u> | <u>44,334</u> | <u>20,873</u> | <u>88</u> | <u>51</u> |

### 3.1 Spatial structure in Bonefish and Queen Conch

Structure runs for *A. vulpes* employing the ALL dataset (=neutral differentiation) revealed scant separation among sample sites (Figure 3). The most highly-supported models for SD2 and SD3 were K=2 and K=3 groupings, both of which depicted GSB (Grand Bahama, south side) as clustereing with EWB (Eleuthera, west side) and ES2 (south and west side of Eleuthera). The second cluster placed GNB (Grand Bahama north side) with EEB (Eleuthera east or Atlantic side) and ES1 (south and west side of Eleuthera) (Figure 3). The K=3 Structure run with SD2 and SD3 datasets was a bit mixed, with GSB and EWB as one group with ES1, ES2, and EEB as a second, and GNB as the third. Subsequent K-groups were decidedly more mixed which rendered them indecipherable.

**FIGURE 3.**
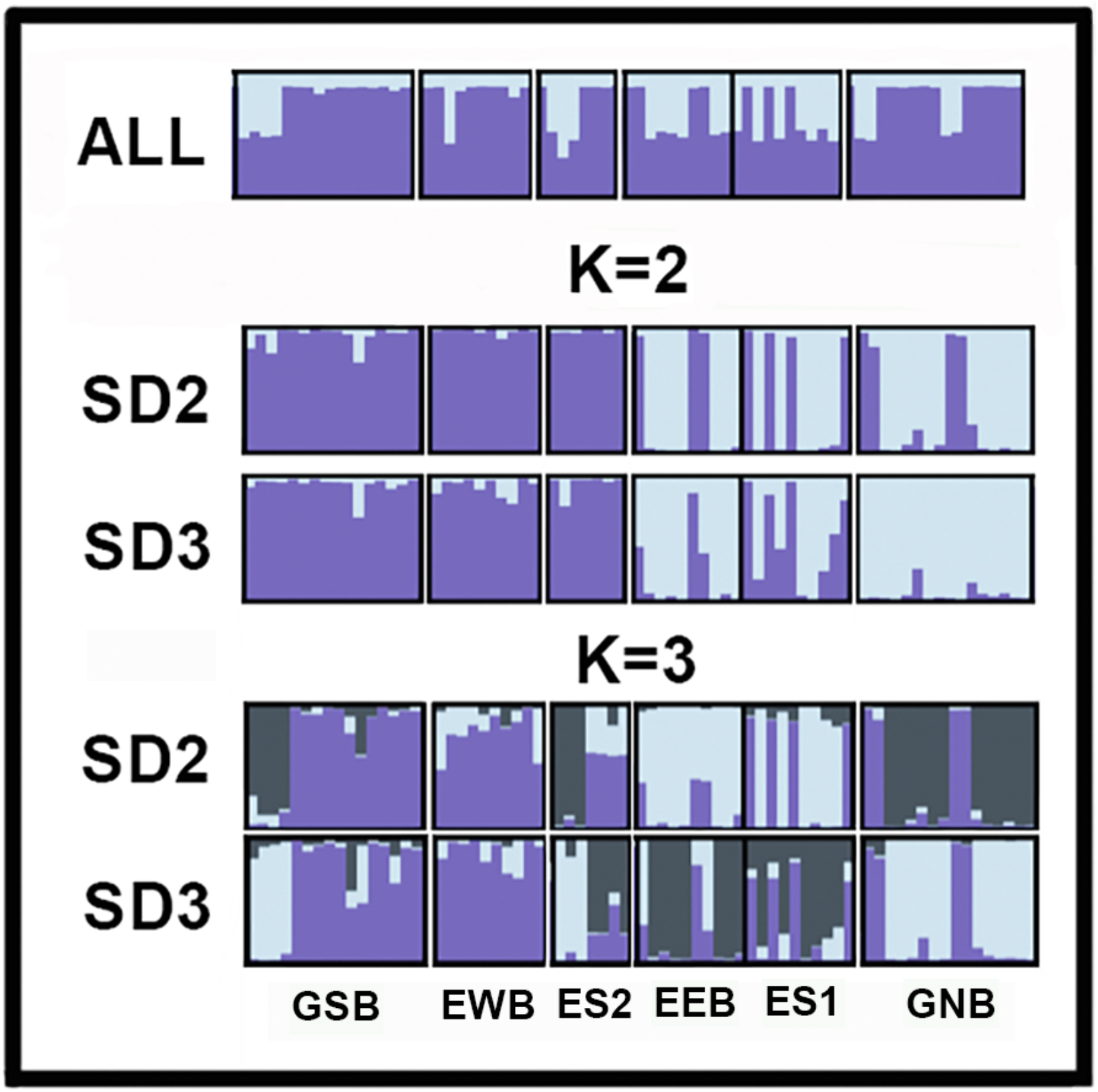
Structure plots for bonefish (*Albula vulpes*) sampled at Grand Bahama and Eleuthera islands, The Bahamas. ALL = Plot employing all single nucleotide polymorphisms (SNPs) (N=25,205); K = 2 represent a 2-group analysis employing: (1) SD2 dataset containing single nucleotide polymorphisms (SNPs) >2 standard deviations (sd) above mean *F*_ST_ (N=123); (2) SD3 dataset containing SNPs >3 sd above mean *F*_ST_ (N=79). K = 3 represents 3-groupl analysis involving the same two datasets. GNB = Grand Bahamas Flats North (Flats or North side); GSB = Grand Bahamas Ocean Side (South or Bahamian side); EWB = Eleuthera Flats Broad Creek (West side); EEB = Eleuthera Ocean Half-Sound (Atlantic Ocean or East side); ES1 = Eleuthera Ocean Plum Creek (Flats or Southwest side); ES2 = Eleuthera Ocean South Side (Flats or Southwest side). Sample site details in Table 1.

In our DAPC analyses, we retained N=45, 25, and 15 PC axes for bonefish (ALL, SD2, and SD3 datasets, respectively), corresponding to 73%, 77%, and 66% of retained variance. In all cases, the number of retained DF axes (N=5) was equal to the number of populations minus one. Clustering of the ALL dataset for bonefish (Figure 4 left) displayed considerable overlap among sites (per Structure plot, Figure 3). Three populations with scant separation (i.e., ES1, ES2 and EEB) were grouped within the center of the graph. Three others were somewhat more peripheral and reflected greater within-group variability (i.e., GNB, GSB, and EWB). The same type of analysis, but employing the SD2 dataset (Figure 4 middle), separated GNB, EWB, EEB, and ES1 without overlap, while grouping GSB with ES2. These relationships were maintained in the DAPC clustering of the SD3 dataset, but with more overlap between the previously separated EEB and ES1, whereas ES2 and GSB maintained their linkage. GNB and EWB also remained distinct (Figure 4 right).

**FIGURE 4.**
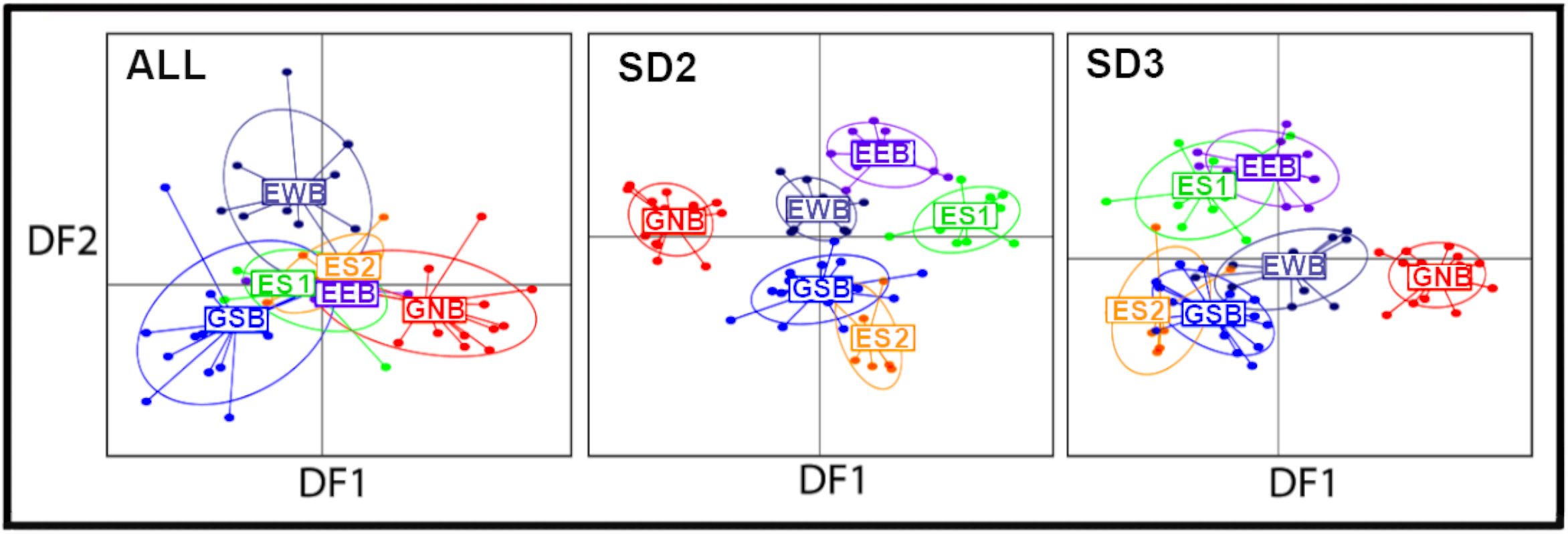
Plots depicting discriminant analysis of principal components (DAPC) for six populations of bonefish (*Albula vulpes*) along discriminant functions one (=DF1) and two (=DF2). ALL = plot utilising all loci (N=25,205); SD2 = plot depicting single nucleotide polymorphisms (SNPs) >2 standard deviations (sd) above mean *F*_ST_ (N=123); SD3 = plot employing SNPs >3 sd above mean *F*_ST_ (N=79). GNB = Grand Bahamas Flats North (Flats or North side); GSB = Grand Bahamas Ocean Side (South or Bahamian side); EWB = Eleuthera Flats Broad Creek (West side); EEB = Eleuthera Ocean Half-Sound (Atlantic Ocean or East side); ES1 = Eleuthera Ocean Plum Creek (Flats or Southwest side); ES2 = Eleuthera Ocean South Side (Flats or Southwest side). Additional details in Table 1.

The Structure analysis of *L. gigas* populations using the ALL dataset revealed little genetic differentiation, with mean *F*_ST_ =0.0129. However both EWC and GSC showed significant separation when both SD2 and SD3 datasets were evaluated (Figure 5). DAPC for conch with cross-validation retained N=15, 10, and 5 PC axes (ALL, SD2, and SD3, datasets, respectively). This yielded 16.7%, 44%, and 50.5% of PC variance. However, there is N=1 DF axis for conch, in that the number of DAPC discriminant functions is N_populations-1. The proportion variance explained by this single axis is irrelevant in that it is 100% by definition. The DAPC clustering of the ALL dataset (Figure 6 left) again showed considerable overlap between the two islands, whereas each separated with considerably less overlap when SD2 was analysed, and became even more distinct based upon SD3 (Figure 6, middle, right).

**FIGURE 5.**
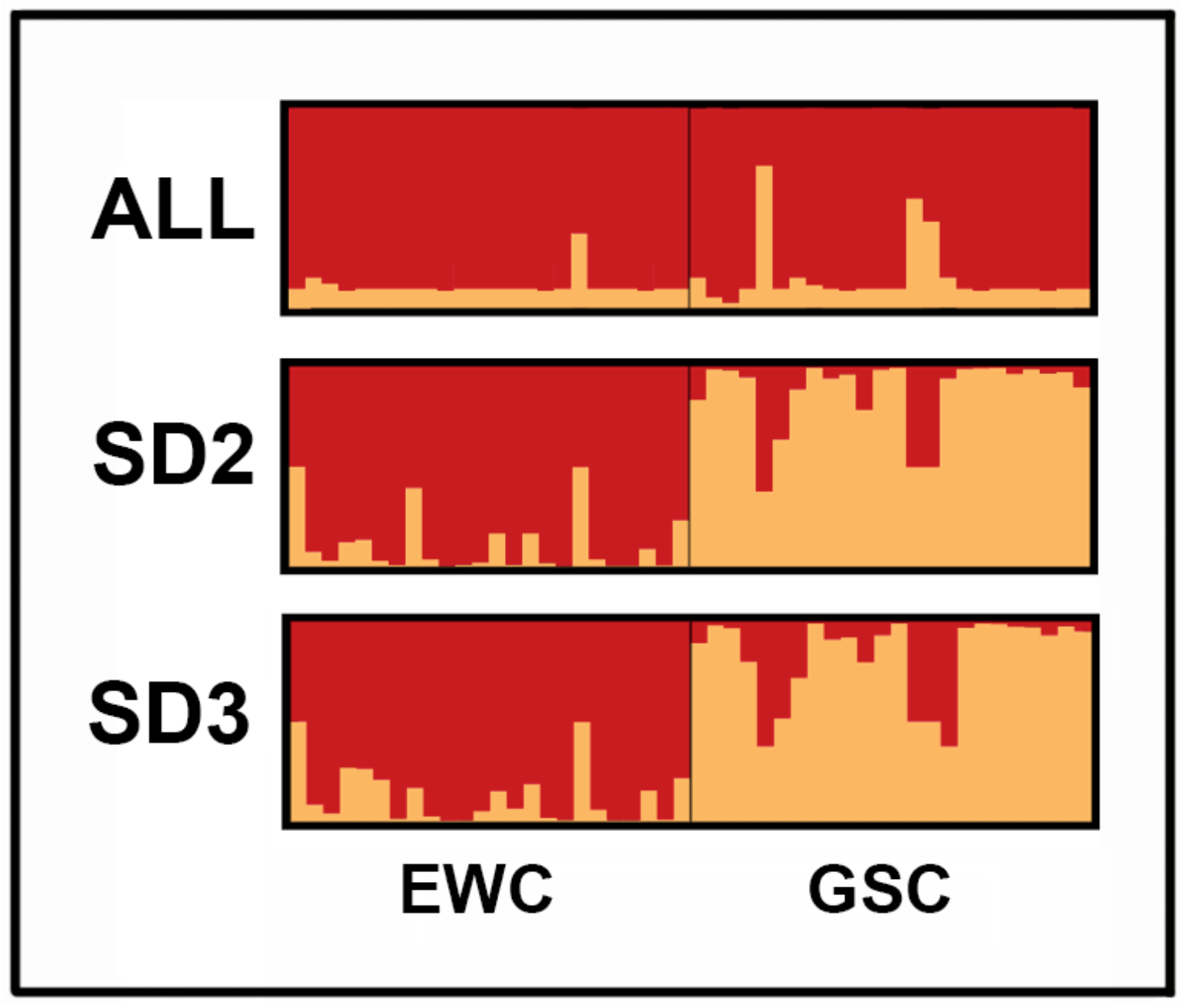
Structure plots for queen conch (*Lobatus gigas*) sampled as Grand Bahama and Eleuthera islands, the Bahamas, with individuals represented by vertical lines. Two groups are depicted [GSC = Grand Bahama South Freeport (Flats or South side); EWC = Eleuthera Flats Schooner Cay 2 (Flats or West side)]; ALL = Plot employing all single nucleotide polymorphisms (SNPs) (N=20,873); SD2 = SNPs >2 standard deviations (sd) above mean *F*_ST_ (N=88); and SD3 = plot depicting SNPs >3 sd above mean *F*_ST_ (N=51).

**FIGURE 6.**
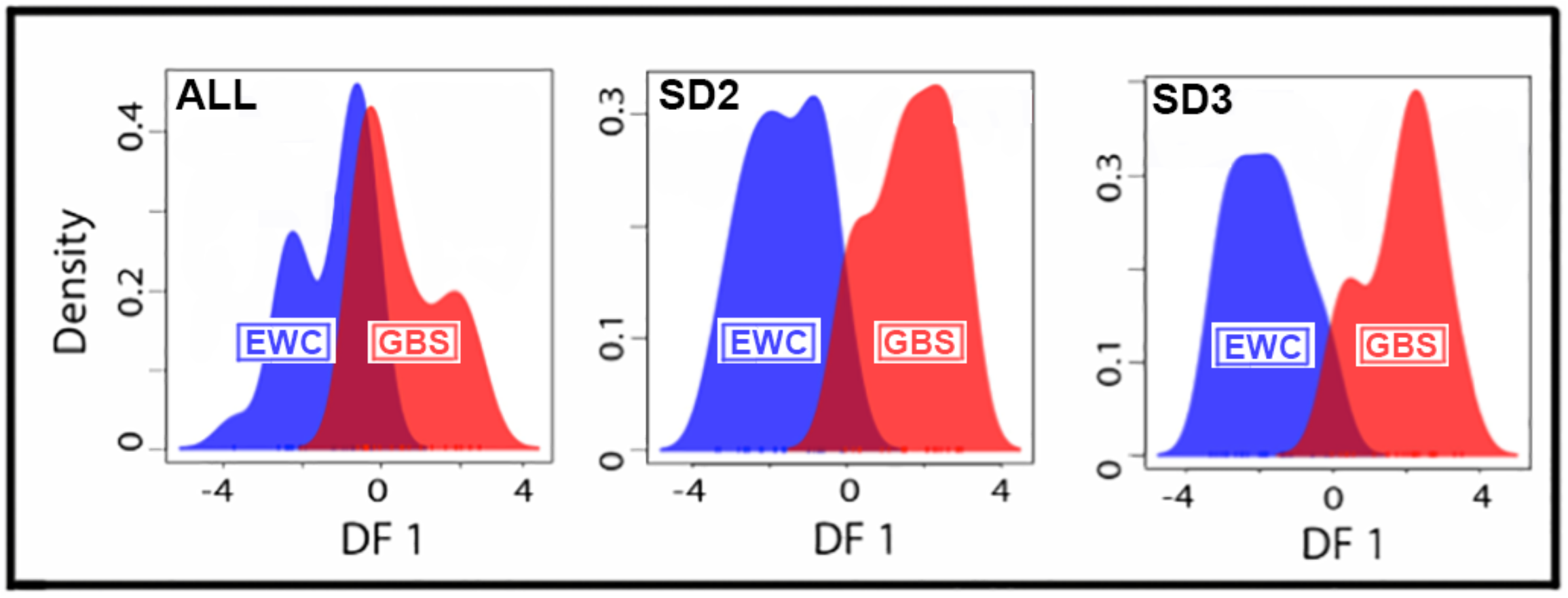
Plots depicting discriminant analysis of principal components (DAPC) for two populations of queen conch (*Lobatus gigas*), with GSC = Grand Bahama South Freeport (Flats or South side) and EWC = Eleuthera Flats Schooner Cay 2 (Flats or West side) plotted along discriminant function one (=DF1). ALL = Plot employing all single nucleotide polymorphisms (SNPs) (N=20,873); SD2 = SNPs >2 standard deviations (sd) above mean *F*_ST_ (N=88); and SD3 = plot depicting SNPs >3 sd above mean *F*_ST_ (N=51).

### 3.2 Asymmetric migration

The BayesAss3 analysis of bonefish revealed highly variable and asymmetric migration rates, with EEB and EWB depicted as source populations, with outward migration rates to both Grand Bahama and flats-side Eleuthera populations. This varied from 8.5-17.9% for EEB and 7.5-14.6% for EWB (Table 3; Figure 7). Notably, the remaining populations lacked any strong signal of outward migration, and all showed negative rates of net emigration (Figure 8), suggesting a sink dynamic. Results for *L. gigas* found very high demographic independence among populations, with ‘self-migration’ (e.g., recruitment) rates estimated at >98% for both Grand Bahama and Eleuthera populations (Table 3).

## 4 DISCUSSION

Populations are frequently classified as to the manner by which they are replenished. In this sense, closed populations are defined as having recruitment via local reproduction, whereas open populations are independent of source (Pinsky et al. 2012). This dichotomy underscores several ecological, evolutionary, and conservation-based research agendas: Exploitation or protection (Yau et al. 2014, Krueck et al. 2017); life history and taxonomy (Mora & Sale 2002, Baco et al. 2016); recovery from disturbance (Bell et al. 2014); and extent to which populations are locally adapted (Jorde et al. 2015, Salles et al. 2016, Segovia et al. 2017). It also has obvious implications for how populations are managed. The advent of genomic techniques has provided considerable resolution, particularly with regard to the estimation of local adaptation (Reisser et al. 2014).

**TABLE 3.** BAYESASS3 results for bonefish (*Albula vulpes*) and queen conch (*Lobatus gigas*) sampled at Grand Bahama and Eleuthera islands. Values are expressed as proportion of individuals expected to be of a given migrant or non-migrant origin. Migration rates along the diagonal are analogous to a self-recruitment rate, with high values representative of demographic independence. Sample site abbreviations per Table 1.

|  | **EWB** | **EEB** | **ES1** | **ES2** | **GNB** | **GSB** | **GNC** | **EWC** |
| --- | --- | --- | --- | --- | --- | --- | --- | --- |
| **EWB** | 0.884 | 0.039 | 0.020 | 0.019 | 0.021 | 0.018 | - | - |
| **EEB** | 0.085 | 0.833 | 0.020 | 0.021 | 0.020 | 0.020 | - | - |
| **ES1** | 0.104 | 0.146 | 0.689 | 0.020 | 0.020 | 0.021 | - | - |
| **ES2** | 0.127 | 0.106 | 0.026 | 0.693 | 0.025 | 0.025 | - | - |
| **GNB** | 0.104 | 0.075 | 0.016 | 0.016 | 0.774 | 0.015 | - | - |
| **GSB** | 0.179 | 0.077 | 0.015 | 0.016 | 0.016 | 0.698 | - | - |
| **GNC** | - | - | - | - | - | - | 0.987 | 0.013 |
| **EWC** | - | - | - | - | - | - | 0.013 | 0.988 |

**FIGURE 7.**
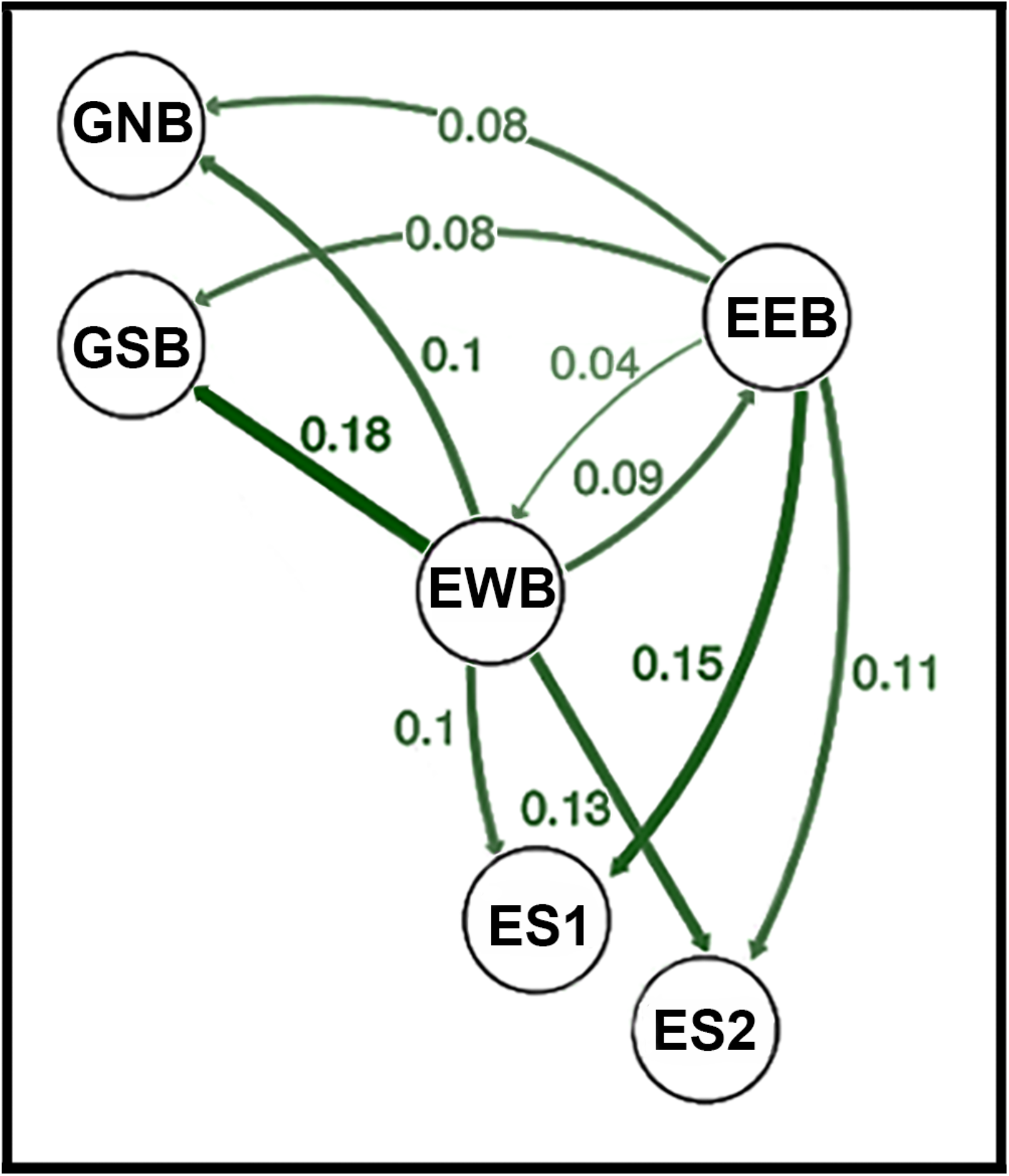
Graph depicting relative migration rates for six populations of bonefish (*Albula vulpes*), where nodes represent populations, and edges are weighted according to the level of direction relative migration rates (*m*), as derived from BA3-SNPs results. All edges below 0.02 were omitted. Full results shown in Table 3. GNB = Grand Bahamas Flats North (Flats or North side); GSB = Grand Bahamas Ocean Side (South or Bahamian side); EWB = Eleuthera Flats Broad Creek (West side); EEB = Eleuthera Ocean Half-Sound (Atlantic Ocean or East side); ES1 = Eleuthera Ocean Plum Creek (Flats or Southwest side); ES2 = Eleuthera Ocean South Side (Flats or Southwest side).

**FIGURE 8.**
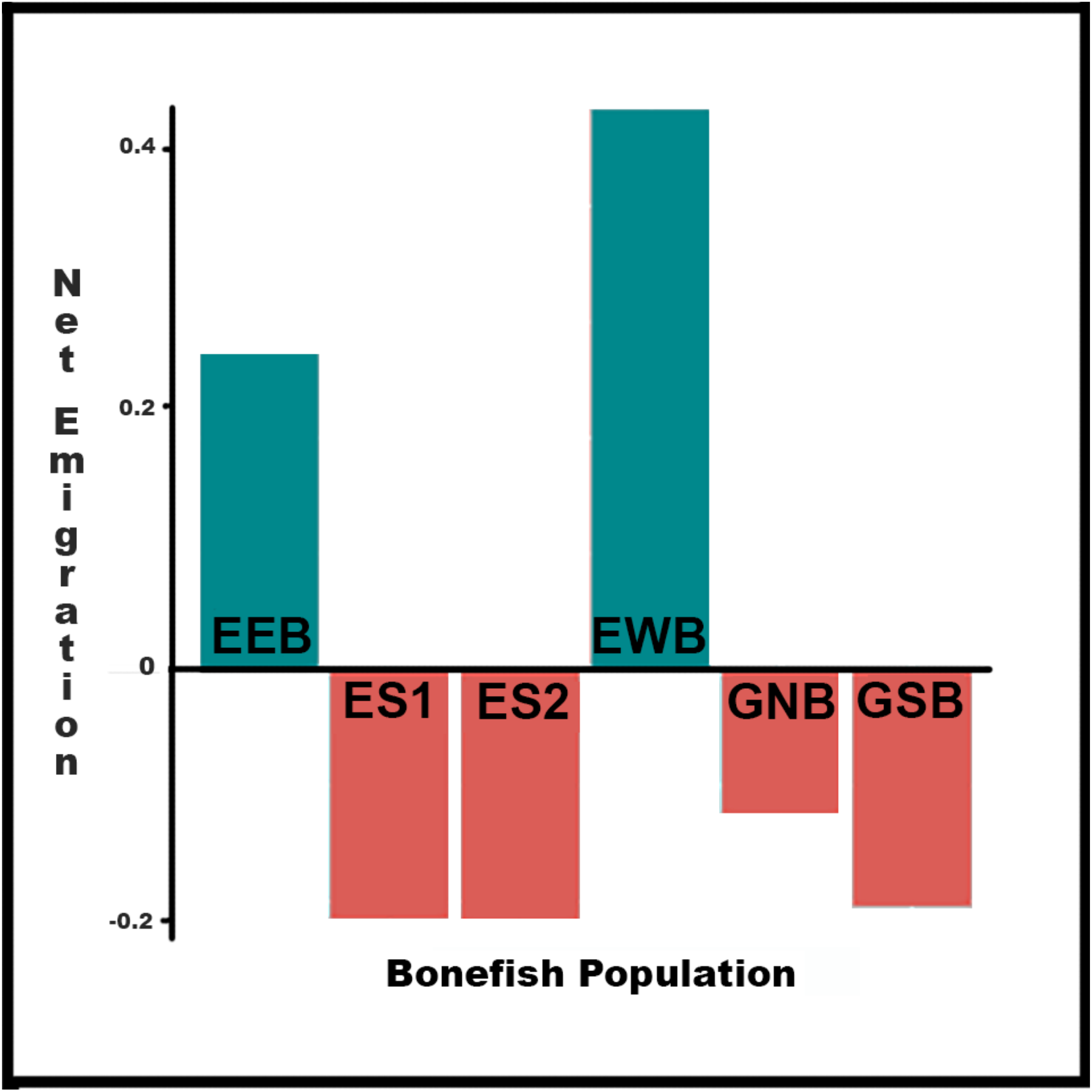
Asymmetric immigration recorded for bonefish (*Albula vulpes*) from Grand Bahama and Eleuthera islands. Values are expressed as net emigration rates (summed emigration - summed immigration) for each population, with positive values indicating ‘source’ populations and negative values indicating ‘sink’ populations. GNB = Grand Bahamas Flats North (Flats or North side); GSB = Grand Bahamas Ocean Side (South or Bahamian side); EWB = Eleuthera Flats Broad Creek (West side); EEB = Eleuthera Ocean Half-Sound (Atlantic Ocean or East side); ES1 = Eleuthera Ocean Plum Creek (Flats or Southwest side); ES2 = Eleuthera Ocean South Side (Flats or Southwest side).

### 4.1 *F_ST_* outliers for bonefish

For non-model organisms (as herein), two approaches are most often utilized: The first identifies loci that display unusually high genetic differentiation among populations (i.e., *F_ST_* outlier approach), whereas the second (i.e., gene-environment association) seeks correlations between the allele frequencies of populations and their local environments. We employed the first approach in this study.

Our analysis failed to group samples of bonefish *Albula vulpes* into geographically congruent groups on either side of the Canyon, suggesting instead a distribution more panmictic in format (per Structure analysis and DAPC plot for the ALL data). Furthermore, Structure plots for K=2 and K=3 groupings (Figure 3) deviated but little. For K=2, both SD2 and SD3 datasets linked the south side of Grand Bahama Island (GSB) with the west side of Eleuthera (EWB) and with one (of two) ES locations (i.e., ES2). This, in turn, suggests a level of connectivity across the Great Bahama Canyon, a conjecture supported by Figure 4. However, the north side of Grand Bahama Island (GNB) grouped with the eastern (Atlantic) side of Eleuthera (EEB) and with the second ES site in the K=2 grouping (i.e., ES1). This relationship separates in K=3, with GSB remaining linked with EWB, whereas EEB and ES2 separate.

These relationships are difficult to interpret biogeographically, particularly given what we know of geomorphic and current patterns in the northeast Basin (Figure 2). A more parsimonious explanation would point to the considerable mobility of adult bonefish superimposed onto a lack of site specificity. Although dispersal pathways for bonefish are relatively unknown in the northeast Basin, their quantification would identify Bahamian populations that serve as sinks for bonefish spawning under investigation at South Abaco Island (Lombardo et al. 2020; Figure 1). In this sense, our results provide reasonable hypotheses.

### 4.2 *F_ST_* outliers for queen conch

A similar analysis did indeed discriminate sessile populations of queen conch *Lobatus gigas* separated by the Great Bahama Canyon (a potentially vicariant barrier). This is aptly demonstrated in Figure 5, where the south side of Grand Bahama Island and the west side of Eleuthera Island are clearly discrete one from another. This condition is reinforced by the DAPC analysis (Figure 6), where distributions for SD2, and SD3 datasets consistently diverge from that depicted for the ALL dataset. *F_ST_* values for these two populations are not significantly different when computed from the ALL data, but are so when compiled from the SD2 and SD3 datasets (Table 3). Thus, while neutral loci demonstrated little differntiation across the study area, a smaller proportion reflected strong differentiation consistent with local adaptation. This scenario has also been manifested in other marine species similarly investigated using SNP analyses (Cure et al. 2017, Thomas et al. 2017).

### 4.3 Potential problems with *F_ST_* outlier analyses

One potential issue with detecting selection via departure from neutrality is that each locus may manifest an independent response that can hinge upon population structure and demography. This is particularly problematic when the average level of differentiation among populations is relatively high (Hoban et al. 2017). However, pairwise *F_ST_* values for bonefish are neither elevated nor significant (Table 3), suggesting this may not apply in our situation. An additional concern is that some loci may demonstrate false signatures of selection, such as when small populations on the leading edge of an expansion contribute disproportionately to observed frequency differences. Again, our study populations are already established and thus the concept of a leading edge is not a viable consideration. Similarly, cryptic hybridization and introgression can invoke confounding effects as well, but again, these are not apparent in our data.

### 4.4 Asymmetrical migration in bonefish and queen conch

The structure of the physical environment can promote differential movements in aquatic organisms, and thus promote a metapopulation structure within a species (i.e., a source– sink dynamic). This in turn can skew estimates of genetic differentiation among populations, and consequently promote erroneous management decisions, particularly when active management is based on an economic model, such as when the organisms are harvested. We employed genetic estimates of population migration to ascertain potential source–sink considerations in our study organisms, and to determine if indeed movements between them were asymmetrical.

Bonefish in our study indeed demonstrated strong signals of source-sink dynamics across islands. The two Eleuthera populations (EEB and EWB) seemingly act as genetic sources, as evidenced by their high net emigration rates derived from BayesAss3 (Figure 8). Their genetic differentiation may suggest the presence of discrete genetic stocks on each side of the island, reflecting either spatial or temporal asynchrony with regards to spawning. Given a tendency for bonefish to move into deeper waters for spawning (Danylchuk et al 2011; Lombardo et al. 2020), the differentiation between the two Eleuthera populations may suggest spawning site-fidelity (potentially within deeper water off Abaco Island; per Lombardo et al. 2020), with increased intermingling subsequently due to the export of larvae to other Bahamian areas. The latter may also explain the cryptic intra-site structure as recorded in the assignment tests of *F_ST_* outlier SNP datasets (Figure 3).

Populations of conch, on the other hand, were highly demographically independent. These results are apparent in both estimates of contemporary migration (BA3-SNPs), and from estimates of population structure using highly-differentiated SNPs.

### 4.5 Dispersal and connectivity within the Caribbean Basin

We hypothesized that the population demographics of our study species would be impacted by the currents passing through the Grand Bahama Canyon. Prior to test, we sought data from other species that might potentially shed light on this conjecture.

Jackson et al. (2014) investigated genetic differentiation among 19 spawning aggregations of Nassau grouper *Epinephelus striatus* in the Basin. The life history of this species parallels that of bonefish, in that spawning aggregations are cued by water temperature and phases of the moon over a 3-month period. As with bonefish, breeding aggregations develop some distance from individual home ranges, with larvae drifting for a similar duration (~40 days) before settling. Four Basin-wide genetic groups were identifed, to include the Bahamas, but with little within-group differentiation, again in parallel with that found for bonefish (per Wallace & Tringali 2016). A major barrier to larval dispersal was the separation of the Basin into eastern and western regions (Jackson et al. 2014, Figure 3). These results implicate the Great Bahama Canyon (Figure 1 top) as a vicariant barrier, similar to what was found herein with conch.

In a second study, Kool et al. (2010) coupled results from a matrix-based projection model with those based upon a bio-physical approach to examine genetic dispersal among coral reef patches within the Basin. Allele frequencies were found to shift along the length of the Bahamas, with the sharpest demarcation occurring at the separation between Grand Bahama Island and the southern Bahamas. This separation was not recognized in the original publication, but indeed juxtaposes with the presence of the Grand Bahama Canyon (Figure 1). Again, the results from Kool et al. (2010) promote the Canyon as a major vicariant feature in the region.

## 5 CONCLUSIONS

An understanding of genetic connectivity within exploited species is a crucial conservation aspect, in that once designated, appropriate geographic units can then be subsequently managed so as to avoid over-exploitation. This includes the establishment of quotas for ‘take,’ as well as the delineation boundaries for marine protected areas (MPAs). Furthermore, an understanding of Basin-wide linkages (or lack thereof) among populations of commercially exploited species (as herein) promotes the realization among numerous stakeholders that fisheries management is a complex and continually evolving geopolitical problem.

Conservation and management practices for targeted species can be greatly enhanced if indeed they are supplemented with adaptive (i.e., ecologically relevant) data (Allendorf 2017). For instance, adaptive-based management units can potentially be defined such that, in high gene flow species, may represent subsets within more traditional (neutral-based) units (Funk et al. 2012). Outlier SNPs could also identify the results of illegal fishing (such as for conch), and also serve to define stock boundaries and potential management units (Milano et al. 2014). For bonefish, it may require the protection of larger areas that can accommodate those fairly precise oceanographic conditions that define where they spawn and which provide the predictable and stable dispersal mechanism required by pelagic larvae.

These aspects cut to the definition of conservation genomics, which is defined (in the narrow sense) as those approaches that differ conceptually and quantitatively from traditional genetics. This, in turn, serves to amplify genome-wide signals of adaptation, and promotes answers for those questions impossible to address using legacy approaches alone (Garner et al. 2016). Importantly, such data can provide genomic evidence for potential heat tolerance (or the lack thereeof), an issue that particularly impacts sessile marine invertebrates (Thomas et al., 2017). Such issues loom large as oceans acidify and their heat profiles expand in lockstep with global climate change (Hurd et al., 2018).

## ACKNOWLEDGMENTS

Funding was provided by the following generous endowments to the University of Arkansas: The Bruker Professorship in Life Sciences (MRD), a Distinguished Doctoral Fellowship (TKC), and the 21^st^ Century Chair in Global Change Biology (MED). We gratefully acknowledge the Cape Eleuthera Institute (CEI) and The Island School for logistical support and field assistance. Sampling was supported under CEI collecting permits. Sampling was supported by grants from Patagonia’s World Trout Initiative, the Charles A. and Anne Morrow Lindbergh Foundation, the Baldwin Foundation, as well as generous personal donations from B. Hallig, T. Rice, and J. Spring. Computational analyses were supported by the Arkansas Economic Development Commission (Arkansas Settlement Proceeds Act of 2000), the Arkansas High Performance Computing Center (AHPCC), and an XSEDE Research Allocation (TG-BIO160065) to access the Jetstream cloud. Sequence data were generated at the Genomics and Cell Characterization Core Facility, University of Oregon/ Eugene. Raw datasets are available via NCBI SRA (accession added upon acceptance).

## AUTHOR’S CONTRIBUTIONS

MRD/ JEC/ DPP/ MED designed the study; JEC/ DPP contributed to fieldwork; MRD/ TKC/ MED contributed to lab work; TKC performed bioinformatic work and statistical analyses; MRD/ TKC/ MED interpreted results and generated figures; MRD/ TKC/ MED drafted the manuscript; All authors gave final approval for publication.

## DATA ACCESSIBILITY

ddRAD SNP loci and scripts can be found at: https://github.com/tkchafin/alb_conch_data

